# Cell2Sentence: Teaching Large Language Models the Language of Biology

**DOI:** 10.1101/2023.09.11.557287

**Authors:** Daniel Levine, Syed Asad Rizvi, Sacha Lévy, Nazreen Pallikkavaliyaveetil, David Zhang, Xingyu Chen, Sina Ghadermarzi, Ruiming Wu, Zihe Zheng, Ivan Vrkic, Anna Zhong, Daphne Raskin, Insu Han, Antonio Henrique de Oliveira Fonseca, Josue Ortega Caro, Amin Karbasi, Rahul M. Dhodapkar, David van Dijk

## Abstract

We introduce Cell2Sentence (C2S), a novel method to directly adapt large language models to a biological context, specifically single-cell transcriptomics. By transforming gene expression data into “cell sentences,” C2S bridges the gap between natural language processing and biology. We demonstrate cell sentences enable the fine-tuning of language models for diverse tasks in biology, including cell generation, complex cell-type annotation, and direct data-driven text generation. Our experiments reveal that GPT-2, when fine-tuned with C2S, can generate biologically valid cells based on cell type inputs, and accurately predict cell types from cell sentences. This illustrates that language models, through C2S fine-tuning, can acquire a significant understanding of single-cell biology while maintaining robust text generation capabilities. C2S offers a flexible, accessible framework to integrate natural language processing with transcriptomics, utilizing existing models and libraries for a wide range of biological applications.

## 1. Introduction

Large language models (LLMs) such as GPT have demonstrated powerful capabilities in natural language processing tasks including question answering, summarization, and text generation (Vaswani et al., 2017; Radford et al., 2018; Devlin et al., 2019; OpenAI, 2023; Touvron et al., 2023; Anil et al., 2023). However, their application to complex fields like biology, especially in single-cell transcriptomics, poses a novel challenge. Traditional methods in this domain, largely reliant on specialized neural network architectures, do not leverage the potential of LLMs’ pretrained knowledge and linguistic understanding.

In this paper, we introduce Cell2Sentence (C2S), a method designed to adapt LLMs for transcriptomics. C2S converts single-cell gene expression data into textual sequences by rank-ordering gene names in descending order of expression levels. This formatting enables LLMs to process and interpret this information (see Figures 1 and 2) while also maintaining the richness and complexity in single-cell data (Figures 3 and 4). Another advantage of C2S is that it takes advantage of the highly optimized and user-friendly open source libraries for transformer models such as Hugging Face (Wolf et al., 2020).

**Figure 1.**
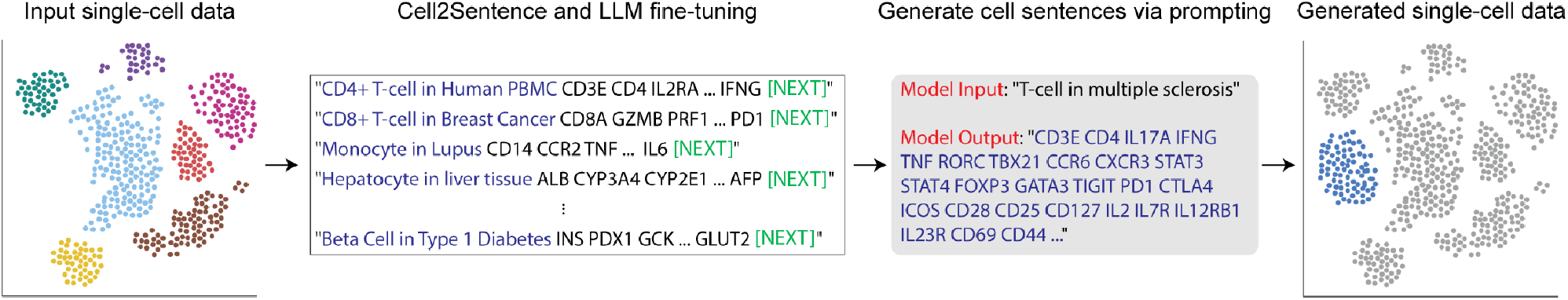
Overview of the Cell2Sentence framework. Input single-cell data, including metadata, are converted into cell sentences for language model fine-tuning. At inference, new cell sentences are generated and can be converted back to gene expression space.

**Figure 2.**
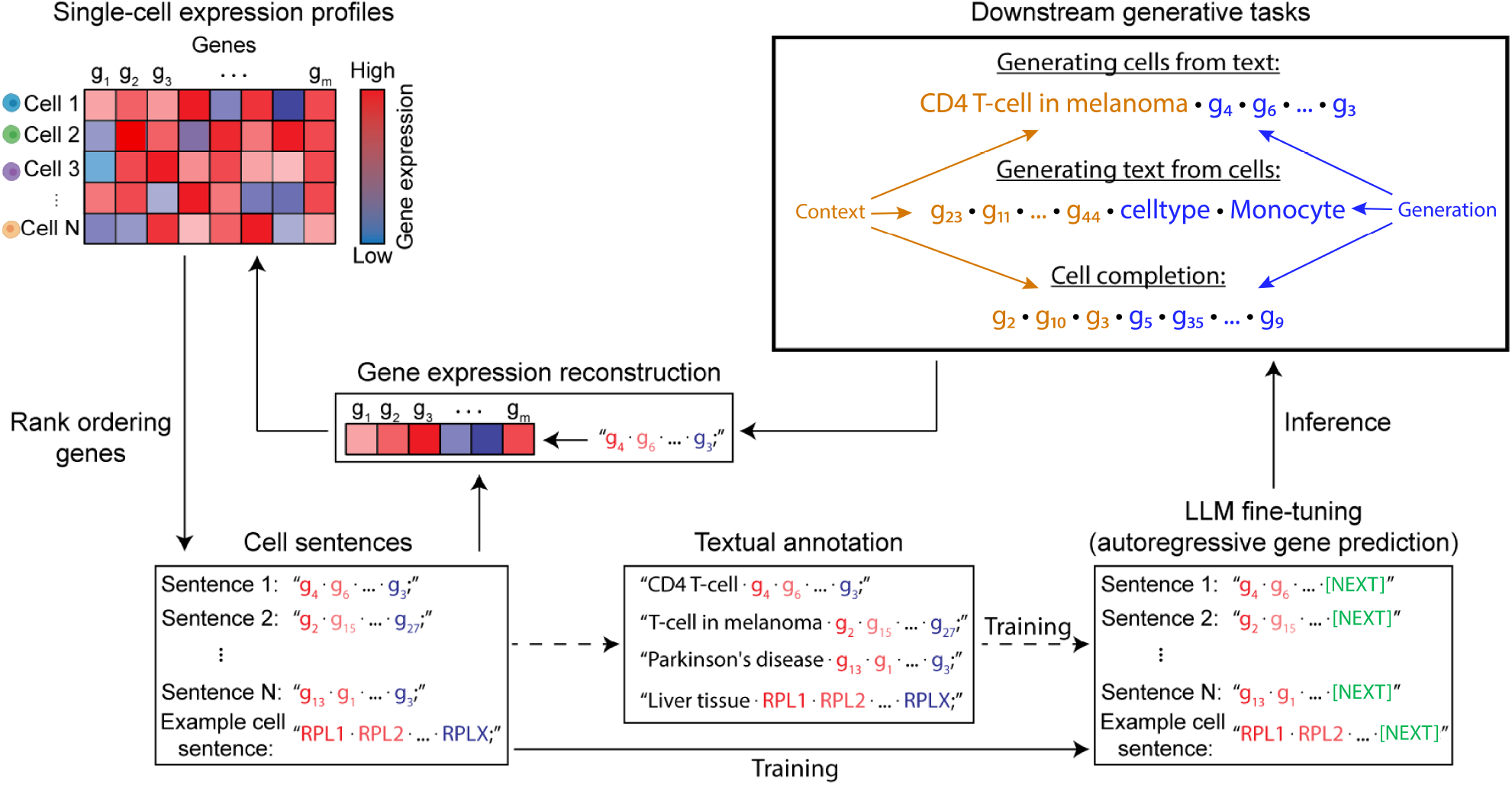
Detailed overview of the Cell2Sentence framework. Single-cell gene expression profiles are transformed into cell sentences through expression rank ordering of gene names. These cell sentences can be annotated with biological metadata, including cell type, tissue, or disease. Subsequently, language models are fine-tuned using the cell sentences. At inference, cell sentences are generated both conditionally (e.g., given a cell or tissue type) and unconditionally, and natural language is produced from text (e.g., cell type label, abstract summary). The generated cell sentences can then be converted back into gene expression profiles.

**Figure 3.**
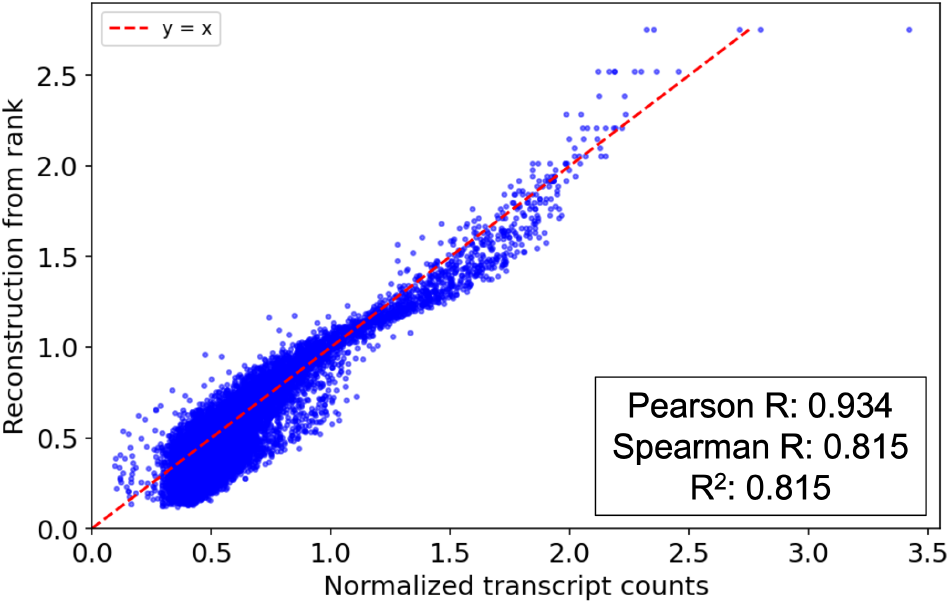
Reconstruction of original gene expression from gene rank by a linear model. Each point in the scatterplot represents one gene sampled from a randomly chosen cell of the PBMC dataset (Domínguez Conde et al., 2022), with 10, 000 genes sampled in total across the dataset for visualization. A linear model accurately reconstructs gene expression from rank across the dataset, with 0.815 *R*^2^ and Spearman correlation against original gene expression, showing that much of the expression information is conserved in the rank-ordered cell sentences.

**Figure 4.**
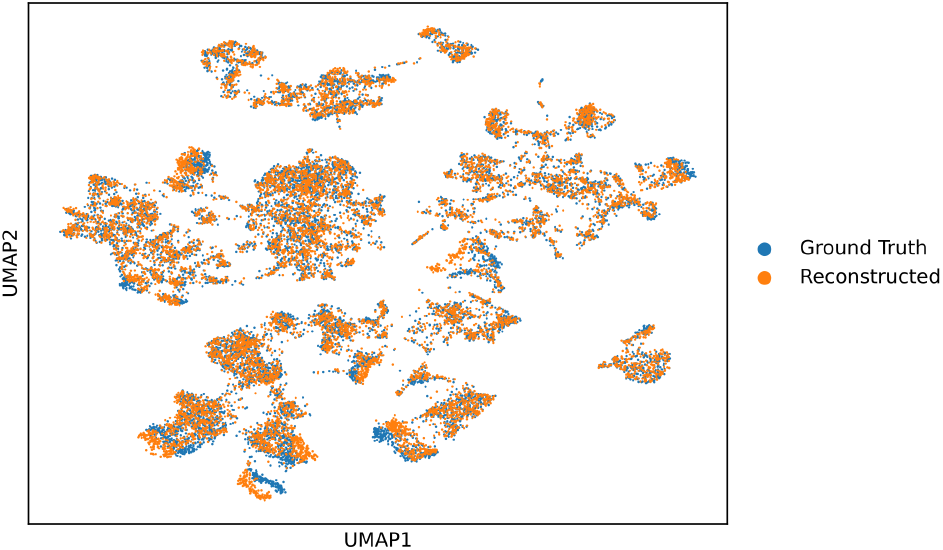
UMAP of ground-truth expression (blue) and reconstructed expression (orange) from cell sentences overlaid. The ground truth cells with their preprocessed expression values from Equation 1 are taken from the PBMC dataset (Domínguez Conde et al., 2022). All 35 cell types with up to 500 cells of each cell type were sampled without replacement for a total of 10350 plotted cells. The reconstructed cells are the ground truth cells whose expression values are reconstructed as follows: first we rank-order the genes and then we reassign the expression values using the learned linear regression parameters from the data. The UMAP qualitatively shows that much of the geometric structure present in the original data remains in the reconstructed data.

This approach not only allows the models to generate biologically relevant cells and predict cell types but also facilitates the generation of descriptive natural language text from single-cell data. This work demonstrates that combining C2S with LLMs significantly enhances their performance in transcriptomic tasks, notably outperforming models trained solely on C2S.

In summary, our key contributions are:

1. Introducing Cell2Sentence, an *effective method* for representing single-cell data as text sequences.

2. Fine-tuning language models on cell sentences to generate and perturb cells, predict combinatorial cell labels, and interpret single-cell data in natural language.

3. Providing a *simple and modular framework* for adapting language models to transcriptomics using popular language model tools and libraries.

In the following sections, we detail the C2S data transformation, model fine-tuning, and evaluate our fine-tuned models on various biological tasks. We conclude with a discussion on the broader implications and future directions for merging natural language processing with transcriptomic data analysis through representation learning. We plan to open-source our software and cell sentence datasets.

## 2. Background and Related Work

### 2.1. Large Language Models

LLMs have transformed natural language processing, demonstrating versatility in tasks ranging from text classification to text generation. Pioneering architectures such as LSTM (Hochreiter & Schmidhuber, 1997), BERT (Devlin et al., 2019), and RoBERTa (Liu et al., 2020) have excelled in text classification, while models like LlaMA-2 (Touvron et al., 2023) and Falcon (Almazrouei et al., 2023) have advanced question answering. Text generation has seen remarkable strides with T5 (Raffel et al., 2020), GPT-3 (Brown et al., 2020), and BART (Lewis et al., 2020). For an extensive exploration of LLMs’ evolution, refer to (Zhao et al., 2023).

### 2.2. Single-Cell Foundation Models

Parallel to LLMs, deep learning in single-cell transcriptomics has progressed significantly. Models like NeuCA (Li & Feng, 2023), ACTINN (Li & Feng, 2023), and scVI (Lopez et al., 2018) have been instrumental in cellular annotation. Tools such as scGen (Lotfollahi et al., 2019) and SAUCIE (Amodio et al., 2019) have addressed batch effect removal, while scVI and DeepImpute (Arisdakessian et al., 2019) have pioneered data imputation. Centralized repositories like GEO (Barrett et al., 2012), CellxGene (Megill et al., 2021), and HCA (Regev et al., 2017) further propelled the field, leading to the development of foundation models like scGPT (Cui et al., 2023b) and Geneformer (Theodoris et al., 2023).

### 2.3. Prompt Fine-Tuning

Prompting, a technique that has been popularized since the introduction of GPT-2 (Radford et al., 2019), has become key to eliciting specific behaviors from LLMs (Lester et al., 2021; Gao et al., 2021; Li & Liang, 2021). The availability of datasets such as Alpaca (Taori et al., 2023a) and tools like Flan (Wei et al.; Longpre et al., 2023; Chung et al., 2022), along with parameter-efficient tuning methods (Hu et al., 2022), has enabled the customization of LLMs for specific tasks. A comprehensive survey of these methods is available in (Liu et al., 2023).

### 2.4. Multimodal Training and Cross-Modality Encoding

Our C2S approach uniquely integrates multimodal learning by transforming single-cell data into text prior to the embedding step (Baltrušaitis et al., 2018; Barua et al., 2023), diverging from traditional schemes that encode modalities separately. This strategy parallels the visual language modeling of (Wu et al., 2007), but uniquely applies to single-cell transcriptomics using modern embedding techniques, marking a novel direction in multimodal machine learning.

## 3. Methods

Cell2Sentence transforms single-cell expression data into sentences of gene names rank ordered by decreasing transcript abundance as shown in Figure 2. While the expression is no longer explicitly contained in the transformed data, we show in Section 3.2 and Figures 3, 4, and 7 that the expression values can be recovered with minimal loss of information. Thus, our method allows for analysis in both rank-order and gene expression formats.

### 3.1. Data transformation

Single-cell RNA sequencing produces transcript count matrices that represent the genetic profiles of individual cells. Most current computational models in single-cell biology handle data in R^*c×n*^, posing scalability challenges with larger datasets. We propose transforming expression matrices into gene sequences as a solution to enable the use of LLMs (Cui et al., 2023a; Hou & Ji, 2023) and other transformer-based architectures (Yang et al., 2022) for single-cell data analysis.

Let *C* denote a matrix with *n* rows and *k* columns corresponding to cells and genes respectively, with *C*_*i*,*j*_ denoting the number of RNA molecules observed for gene *j* in cell

*i*. We filter cells with fewer than 200 genes expressed and genes expressed in less than 200 cells. Cell-wise quality control metrics are then calculated based on mitochondrial gene counts using the Scanpy Python library (Wolf et al., 2018). Cells with over 2500 counts or more than 20% mitochondrial transcript counts are excluded. The count matrix is then row-normalized (summing to 10,000) and log-normalized (Haque et al., 2017), yielding the preprocessed transcript count matrix *C*^*′*^. We summarize this normalization step as:

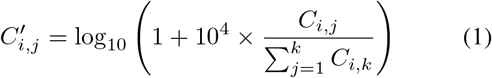

We denote the rank-order transformation applied on *C*^*′*^ as *S*, and the sequence of gene names resulting from *S*(*C*_*i*_) as cell sentence *s*_*i*_ for each cell *i* in the preprocessed count matrix. In practice, we apply the preprocessing and rank-order transformation *S* on each individual single-cell dataset, providing a flexible process for converting traditional single-cell gene expression count matrices to cell sentences.

While genes are not intrinsically ordered in transcript matrices, their expression patterns have been shown to follow inverse-rank frequency patterns (Furusawa & Kaneko, 2003; Qiu et al., 2013), thus establishing a steady relationship between a gene’s expression level within a cell and its rank among the genes expressed in that cell. We model this inverse-rank relationship with a log-linear distribution and approximate it in log-log space using a linear regression (Dhodapkar, 2022).

Given single-cell dataset which underwent rank-order transformation *S*, let *r*_*i*_ denote the log of the rank of gene *i* in *C*^*′*^, and *e*_*i*_ the original expression of gene *i*. We fit a linear model to predict *e*_*i*_ from *r*_*i*_ during the initial conversion to cell sentence format, resulting in a fitted slope and intercept value which are saved for each converted dataset (see Figures 7). The linear model has form *e*_*i*_ = *a*_*d*_ *× r*_*i*_ + *b*_*i*_, given dataset *d* and *{a*_*d*_, *b*_*d*_*}* ∈ *ℝ*^2^.

We postprocess generated cell sentences by ignoring invalid gene names and averaging the rank of duplicate genes. The fitted linear model is then applied to the log-rank of generated genes to convert back to expression. Any gene not present in a cell sentence is considered to have zero expression. We define the average rank of a generated gene 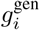 belonging to the set of unique genes *G*_*U*_ ⊆ *S* as follows:

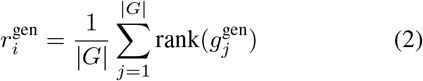

where 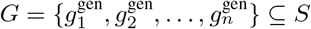 is the set of duplicate generated genes for 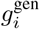, and 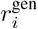 denotes the average rank of gene 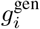 in the generated cell sentence. This yields the following formulation for expression value vector for the generated cell

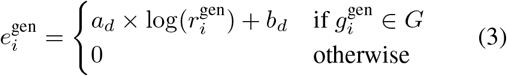

In practice, we consider a global dictionary of all gene names seen in single-cell datasets, which dictates the size of the resulting gene expression vector of the cell.

Next, we consider the robustness of the C2S transformation when converting cells to sentences and reverting back to expression.

### 3.2. Transformation robustness

We find that transforming expression data to cell sentences is a robust and reversible operation, and cells converted to text and back to expression incur minimal information loss, with over 81% of the variation in the gene expression being captured by the linear regression on an immune tissue dataset (see Figures 3 and 4). Relationships between normalized gene expression and rank on a variety of human tissue datasets are shown in Figures 6 and 7 (see Appendix A). As the linear model only requires the log-rank of a gene to approximate its expression level, any gene sequence can be converted to expression, including generated cell sentences.

**Figure 5.**
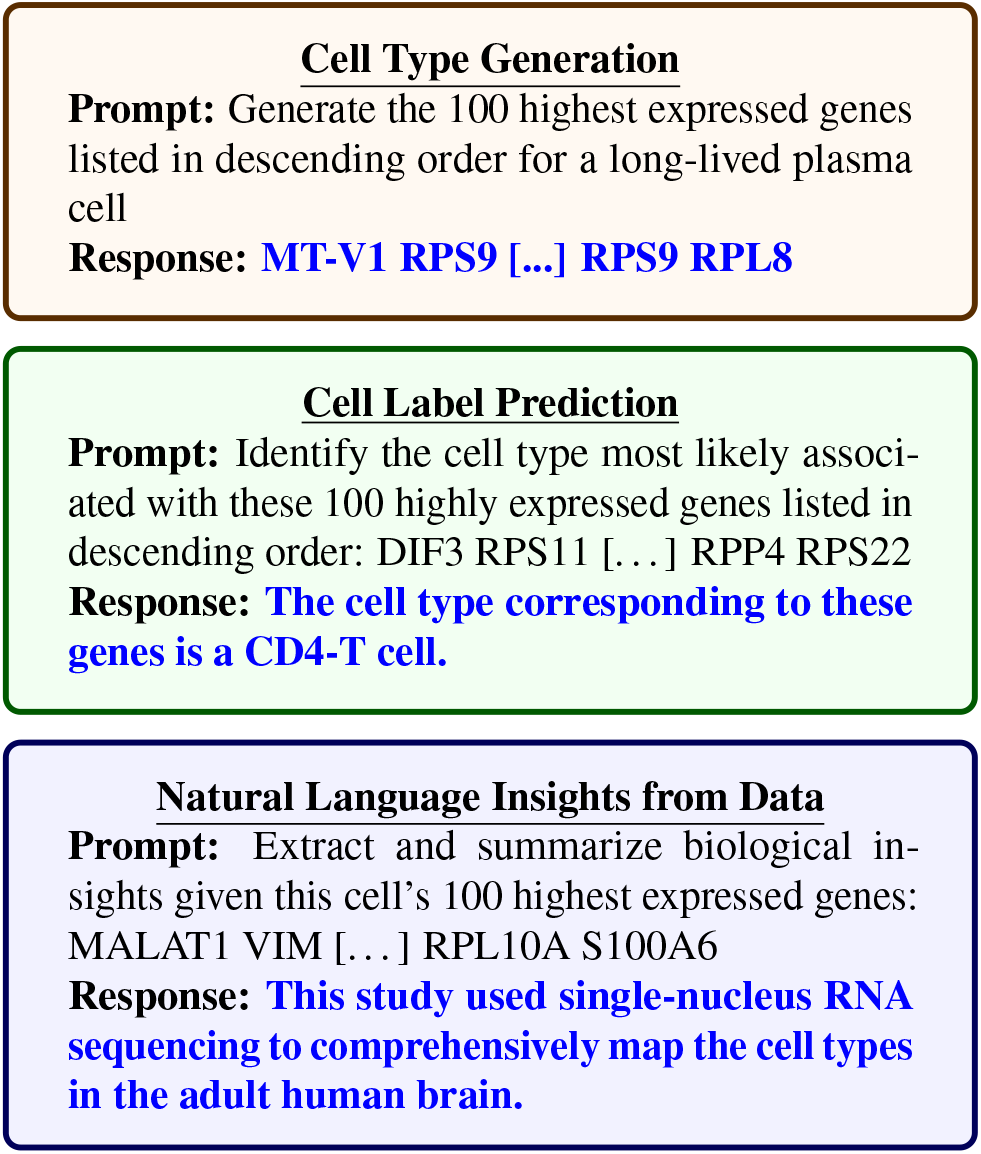
Example of Cell2Sentence prompts and responses for generating cell sentences from text, predicting complex natural language labels, and generating biological insight from a single cell sentence.

**Figure 6.**
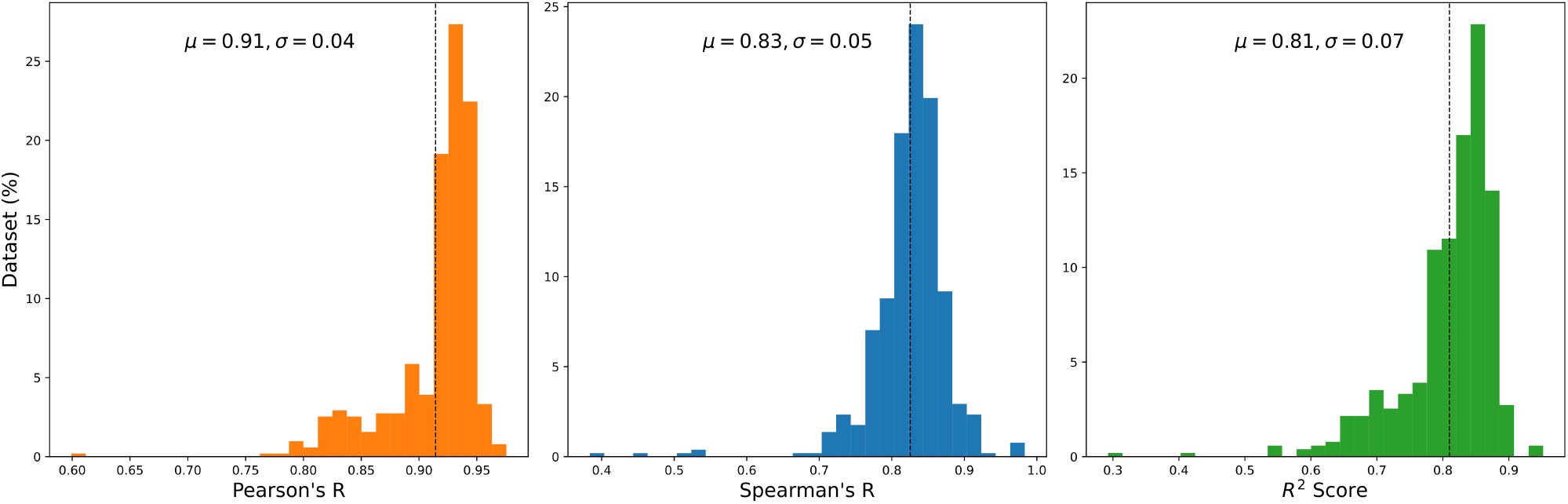
Distributions of Peason’s *R*, Spearman’s *R* and *R*^2^ statistics for linear expression reconstruction from log-rank using across 127 distinct scRNA-seq datasets. Both mean and standard deviation are shown for each score, with the mean represented by a dashed line on the x-axis.

**Figure 7.**
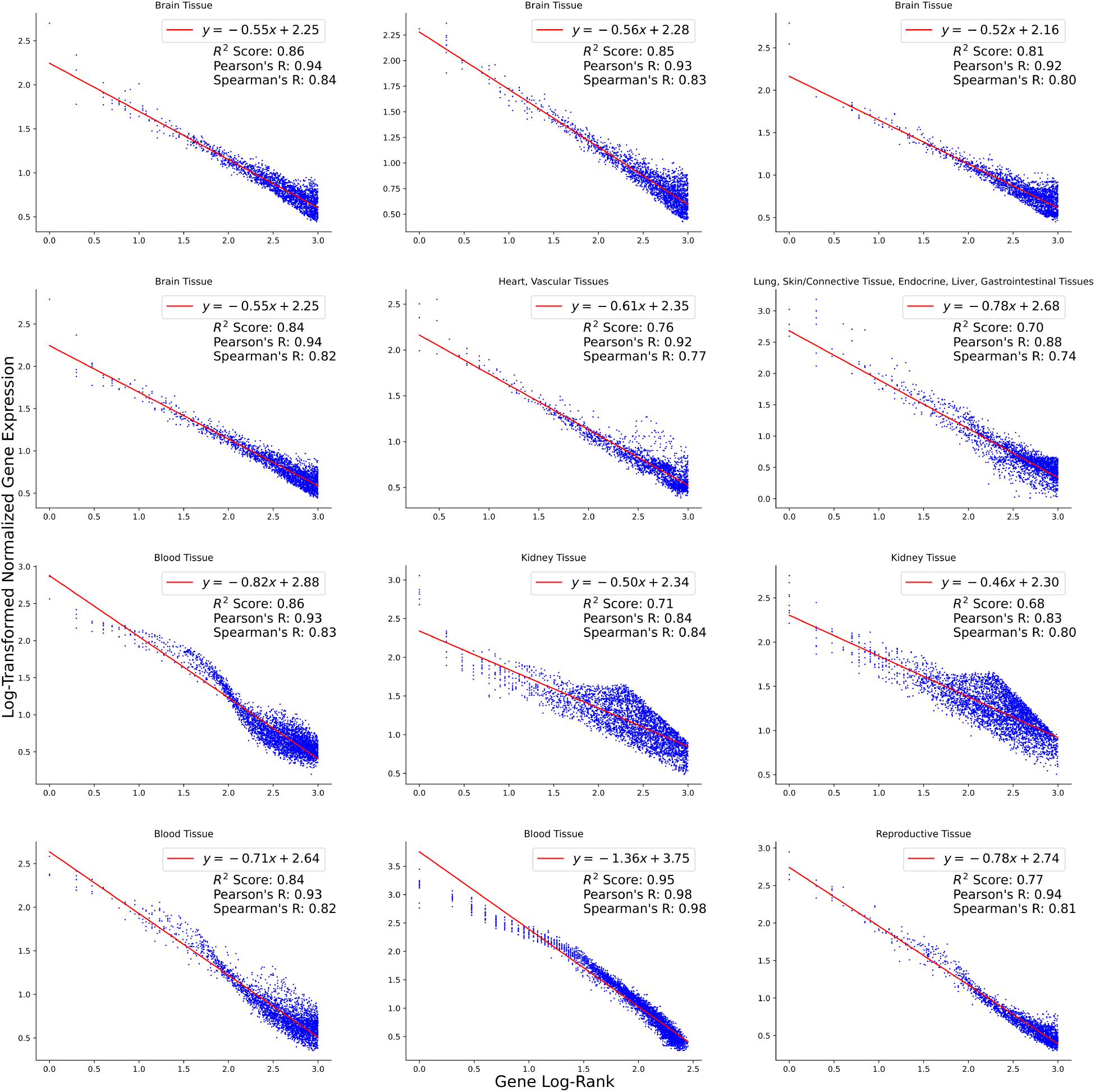
Scatter plots showing the log-linear relationship between gene expression and rank across 12 diverse scRNA-seq datasets. The red line shows the fitted linear model, with Pearson’s *R*, Spearman’s *R*, and *R*^2^ quantifying goodness of fit. The high correlation values demonstrate that gene rank encodes expression in a consistent, reversible way across datasets. This enables translating between the text domain of cell sentences and original gene expression.

### 3.3. Tasks

The objective of this work is to train language models to generate single-cell data and derive biological insights from single-cells in natural language. We summarize the tasks implemented in our experiments (see Section 4):

- Generate cell sentences: C2S models are trained to generate gene sequences from prompts, optionally conditioned on additional metadata.
- Predict cell labels: biological experiments often involve combinatorial labels (e.g., patient and sample metadata) which C2S models can learn to predict from cell sentences directly in text.
- Derive natural language insights: C2S analyzes single-cell data to extract high-level information on expression dynamics by pairing relevant natural language with cell sentences.

Examples of prompts and responses for each of these tasks are shown in Figure 5. For the cell generation and natural language tasks, the models were trained using the standard causal language modeling loss:

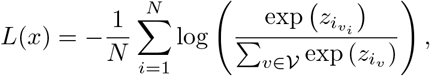

where *x* = *{x*_0_, *x*_1_, …, *x*_*N*_ *}* is the input sentence with *N* + 1 tokens, the model output at position *i* is *z*_*i*_ *∈* R^|*V*|^, and 𝒱 is the vocabulary of the language model with *v*_*i*_ the ground truth token at position *i* in the tokenization of *x*. The loss is averaged over each batch.

## 4. Experiments

In this section we benchmark Cell2Sentence on conditional cell generation, combinatorial cell label prediction, and abstract generation from cell sentences. These experiments are conducted using models fine-tuned either with an immune tissue dataset (Domínguez Conde et al., 2022) or with a large scale multi-tissue dataset (Megill et al., 2021). In both cases, we fine-tune GPT-2 (small, medium and large) (Radford et al., 2019) using cell sentences truncated to 100 genes due to resource constraints. We also fine-tune Pythia-160m (Biderman et al., 2023), which is based on the GPT-NeoX architecture (Black et al., 2022) and uses rotary embeddings (Su et al., 2024). For the latter, we set the model’s maximum input sequence length to 9200 tokens during fine-tuning, permitting manipulation of full cell sentences. We structure our experiments by first fine-tuning language models on large single-cell datasets, and optionally continuing training for downstream evaluation tasks (see Appendix B.2).

### 4.1. Fine-Tuning Datasets

We focus our experiments on three datasets with extensive natural language metadata and labels, allowing to leverage the capabilities of base models.

#### Immune tissue

(Domínguez Conde et al., 2022) proposes a large human immune tissue single-cell dataset with cell type annotations. After transformation, we obtain 273,502 cell sentences, each paired with one of 35 cell type labels. We hold out 20% of cell sentences for validation (10%) and testing (10%). We derive three tasks for this dataset:

- Unconditional cell generation (generate a random cell without any specified cell label).
- Cell type generation (generate a cell sentence given a specific cell type).
- Cell type prediction (predict the cell type in natural language given a cell sentence prompt).

We create 20 prompt templates to embed cell sentences and labels in natural language.

#### Cytokine stimulation

(Dong et al., 2023) is a single-cell dataset that applies 9 cytokine stimulation combinations to immune tissue and with 2 different exposures. Cells are split into 7 cell types for a total of 140 combinatorial labels including unstimulated control cells. This dataset is used in 2 tasks:

- Perturbed cell generation (generate a perturbation given only the labels in text format).
- Cell label classification (classify the cell type, perturbation, and exposure based on the input cell).

In the perturbed cell generation task, 10 out of 140 combinatorial labels were held out during training to be used at test time. For cell label classification, all combinations were used during training, but a limited amount of data was used during training.

#### Multi-tissue

(Megill et al., 2021) provides access to hundreds of human and mouse single-cell datasets. We select 99 human single-cell datasets and convert each of them to cell sentences (see Section 3), yielding a total of 37M cells (including 19 held out studies, representing 2.7% of all cell sentences). Every cell sentence is paired with a tissue label derived from the study’s metadata (e.g., the tissue label for a dataset containing “Brain” and “Liver” cells will be “Brain, Liver”). We find a total of 11 unique tissues, and 42 unique tissue combinations across this dataset. Additionally, we generate synthetic abstract summaries to augment original abstracts for the multi-tissue dataset (see Appendix B.1). We derive the following five tasks for this multi-tissue dataset:

- Tissue type prediction and conditional generation (similarly to the previous cell type prediction and conditional generation prompts).
- Generate abstract summaries from cell sentences (prompted with a cell sentence, the model generates an abstract summary for the corresponding study).
- Generate a cell sentence given an abstract summary (similar to generating from tissue or cell type, but in-stead leveraging natural language from abstract summaries).

We also include an unconditional generation prompt template, where models are only prompted to generate a cell sentence.

### 4.2. Experiment 1: conditional cell generation

#### Objective

Accurate generation of different cell types is crucial for generative approaches on single-cell data, as it enables downstream analysis. Our aim is to establish the quality of several cell generation methods by using distributional, correlation, and classification metrics. We show that C2S models outperform state-of-the-art baselines in generating synthetic single-cell data given cell type and perturbation conditions.

#### Methodology

- **Cell type generation** We train C2S on the immune tissue dataset from (Domínguez Conde et al., 2022). Cell sentences are appended to text prompts indicating its cell type in order for the model to learn cell type generation (Figure 5). Our approach was compared against several established generative single-cell methods, including scVI, scGen, scDiffusion, and scGPT (Lopez et al., 2018; Lotfollahi et al., 2019; Luo et al., 2024; Cui et al., 2023a). For scGen, we randomly split our training data in half and artificially set one group as *control* and the other as *stimulation*, and we randomly pair cells of the same cell type. scGPT’s unique conditional generation method was described in (Cui et al., 2023a), but no code for this method was made publicly available as of this writing. We mimic their method by adding an embedding layer for cell type labels and add the embedding to all tokens while generating random samples of 1000 genes until 36000 genes are generated. A held out test dataset of 500 cells per cell type (17,500 cells in total) with 36,503 genes was used to evaluate all models. We run our experiments 5 times using 5 different samples of our test dataset and report the mean score and standard deviation. Further details on training and data are supplied in Appendices B.3 C.0.3 and Table 6.
- **Perturbed cell generation** We use the single-cell cytokine stimulation dataset from (Dong et al., 2023). The prompts are constructed similarly to the cell type generation task except with additional cytokine stimulation and exposure labels for a total of 3 labels per cell sentence. The C2S and scGen models were trained on all 21710 genes remaining after standard filtering using the scanpy library (Wolf et al., 2018). scGPT was trained on only the 5000 highly variable genes reported in Table 2. Further details on methodology can be found in Appendices B.3.

**Table 1.** Results on immune cell conditional generation quality. For each measure we sample 500 cells with replacement from each cell type in our held out immune dataset for comparison. The *k*-nearest neighbors (*k*-NN) classifier is fit on the held out immune cells with their cell types used as labels. The true label of a generated cell is the cell type used for its conditional generation. Gromov-Wasserstein (GW) distance is measured between all generated and all held out cells. Our full cell generation C2S model outperforms all models.

| MODEL | $k$-NN (↑) | | | | GW (↓) |
| --- | --- | --- | --- | --- | --- |
|  | 3 | 5 | 10 | 25 |  |
| SCGEN | $0.2376 \pm 0.0112$ | $0.2330 \pm 0.0093$ | $0.2377 \pm 0.0053$ | $0.2335 \pm 0.0041$ | $315.9505 \pm 1.2431$ |
| SCVI | $0.2436 \pm 0.0062$ | $0.2400 \pm 0.0064$ | $0.2425 \pm 0.0034$ | $0.2348 \pm 0.0032$ | $302.1285 \pm 0.9338$ |
| SCDIFFUSION | $0.2335 \pm 0.0125$ | $0.2288 \pm 0.0111$ | $0.2368 \pm 0.0067$ | $0.2306 \pm 0.0049$ | $72.0208 \pm 0.3937$ |
| SCGPT | $0.1838 \pm 0.0086$ | $0.1788 \pm 0.0169$ | $0.1811 \pm 0.0149$ | $0.1882 \pm 0.0071$ | $2989.8066 \pm 4.9229$ |
| C2S (PYTHIA-160M) | **$0.2588 \pm 0.0061$** | **$0.2565 \pm 0.0060$** | **$0.2746 \pm 0.0073$** | **$0.2715 \pm 0.0070$** | **$54.3040 \pm 0.3410$** |

**Table 2.** Predicting perturbation effects on unseen conditions. We use a dataset comprising combinatorial cytokine stimulation of immune cells (Dong et al., 2023). The Pearson R and Spearman R values are computed using the mean expression vectors in the unseen test dataset and the corresponding mean expression vectors generated by the model. The top 5000 highly variable genes are selected based on the training dataset, and the top 20 most differentially expressed genes between exposures of the same cell type and perturbation are computed from those 5000 highly variable genes. The Δ symbol indicates the correlations based on differencing with the mean expression vector of the opposite exposure in the training dataset. The conditioning labels are combinatorial, consisting of triples of cell type, cytokine stimulation, and exposure. There are 140 possible combinations in total, and the models were tasked with generating 10 cell type/perturbation combinations with different exposures from those seen during training. Our C2S-trained Pythia-160m model shows superior performance in generating unseen exposures of perturbations compared to SOTA perturbation methods scGEN and scGPT.

|  | MODEL | PEARSON R | TOP-20 DE PEARSON R | SPEARMAN R | TOP-20 DE SPEARMAN R |
| --- | --- | --- | --- | --- | --- |
| | SCGEN | $0.6805 \pm 0.0075$ | $0.7187 \pm 0.0054$ | $0.5654 \pm 0.0025$ | $0.6256 \pm 0.0074$ |
| | SCGPT | $0.0041 \pm 0.0018$ | $0.1299 \pm 0.0495$ | $-0.0002 \pm 0.0040$ | $0.1440 \pm 0.0996$ |
|  | C2S (PYTHIA-160M) | **$0.9241 \pm 0.0002$** | **$0.9734 \pm 0.0007$** | **$0.6210 \pm 0.0005$** | **$0.9752 \pm 0.0016$** |
| $\Delta$ | SCGEN | $0.3871 \pm 0.0060$ | **$0.3743 \pm 0.0167$** | $0.2586 \pm 0.0030$ | $0.3999 \pm 0.0103$ |
| | SCGPT | $0.0980 \pm 0.0001$ | $0.0213 \pm 0.0001$ | $0.0689 \pm 0.0019$ | $0.0441 \pm 0.0006$ |
|  | C2S (PYTHIA-160M) | **$0.4829 \pm 0.0008$** | **$0.3789 \pm 0.0042$** | **$0.2895 \pm 0.0009$** | **$0.4528 \pm 0.0092$** |

**Table 3.** Experimental results on downstream cell label classification. Cell labels are composed of multiple combinatorial metadata parts, including cell type, perturbations, and dosage information. Accuracy and area under ROC curve is computed on model predictions versus ground truth combinatorial labels, with partial credit given for partial misclassifications.

|  | MODEL | CYTOKINE STIMULATION |  | L1000 |  | GTEx |  |
| --- | --- | --- | --- | --- | --- | --- | --- |
|  |  | ACC | AUROC | ACC | AUROC | ACC | AUROC |
| PARTIAL LABEL | k-NN CLASSIFIER | 0.462 ± 0.0047 | 0.550 ± 0.0064 | 0.592 ± 0.0054 | 0.740 ± 0.0036 | 0.492 ± 0.0047 | 0.662 ± 0.0035 |
| | XGBOOST | 0.515 ± 0.0175 | 0.631 ± 0.0202 | 0.389 ± 0.0066 | 0.534 ± 0.0044 | 0.482 ± 0.0127 | 0.633 ± 0.0120 |
| | GENEFORMER | 0.600 ± 0.0170 | 0.722 ± 0.0145 | 0.419 ± 0.0153 | 0.632 ± 0.0181 | 0.500 ± 0.0013 | 0.649 ± 0.0025 |
| | scGPT | 0.419 ± 0.0001 | 0.500 ± 0.0000 | 0.334 ± 0.0076 | 0.500 ± 0.0000 | 0.270 ± 0.0522 | 0.500 ± 0.0000 |
|  | C2S (GPT-2 LARGE) | **0.639 ± 0.0049** | **0.767 ± 0.0049** | **0.631 ± 0.0031** | **0.768 ± 0.0021** | **0.575 ± 0.0035** | **0.713 ± 0.0014** |
| FULL LABEL | k-NN CLASSIFIER | 0.060 ± 0.0044 | 0.509 ± 0.0013 | 0.168 ± 0.0041 | 0.583 ± 0.0020 | 0.112 ± 0.0027 | 0.500 ± 0.0001 |
| | XGBOOST | 0.087 ± 0.0088 | 0.528 ± 0.0050 | 0.124 ± 0.0032 | 0.556 ± 0.0018 | 0.107 ± 0.0080 | 0.500 ± 0.0003 |
| | GENEFORMER | 0.134 ± 0.0161 | 0.555 ± 0.0085 | 0.102 ± 0.0288 | 0.549 ± 0.0146 | 0.123 ± 0.0005 | 0.500 ± 0.0000 |
| | scGPT | 0.049 ± 0.0002 | 0.500 ± 0.0000 | 0.026 ± 0.0001 | 0.500 ± 0.0000 | 0.006 ± 0.0028 | 0.500 ± 0.0000 |
|  | C2S (GPT-2 LARGE) | **0.149 ± 0.0057** | **0.564 ± 0.0030** | **0.202 ± 0.0059** | **0.600 ± 0.0029** | **0.152 ± 0.0062** | **0.574 ± 0.0032** |

**Table 4.**
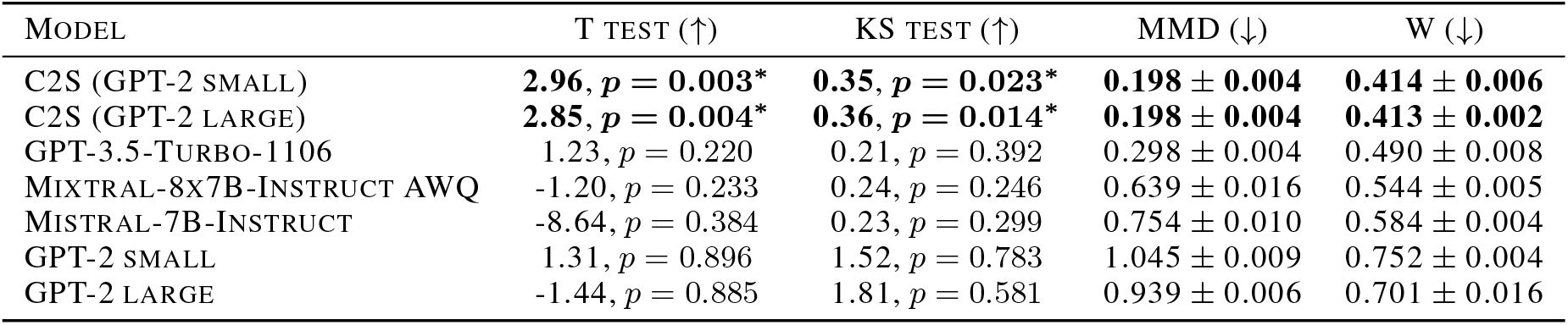
Experimental results on abstract summary generation. This Table displays the outcomes of statistical analyses (T-test, KS test) to evaluate the mean cosine similarities between embeddings of generated abstracts and their respective original abstracts (where *i* = *j*) as well as with different original abstracts (where *i≠ j*). Additionally, it details the Maximum Mean Discrepancy (MMD) and Wasserstein distance (W) comparisons of the embeddings from generated abstracts against those of original abstracts. The results indicate that C2S significantly surpasses baseline methods by achieving a higher correlation with the original abstracts’ embeddings. Furthermore, the embeddings from C2S-generated summaries exhibit closer alignment to those of the original abstracts compared to baseline approaches.

**Table 5.**
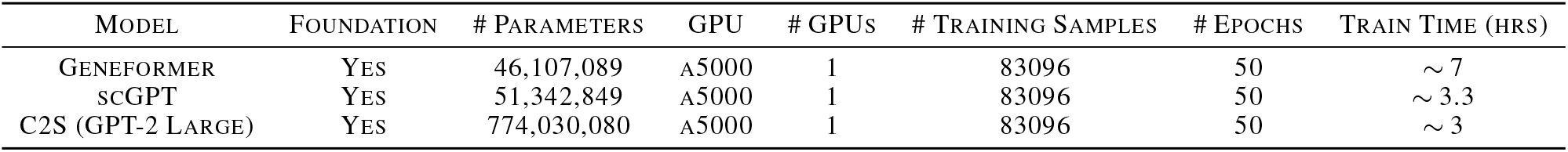
Comparison of required compute on the L1000 dataset (Subramanian et al., 2017). All reported numbers are model checkpoints used in experiments.

**Table 6.**
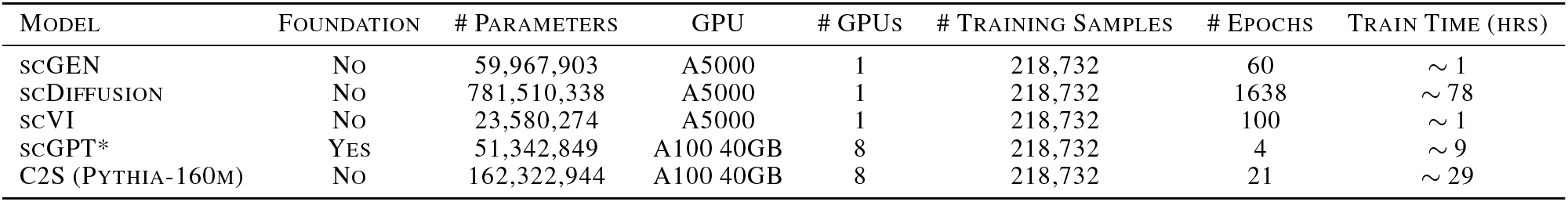
Comparison of required compute on the immune tissue dataset (Conde et al., 2022). All reported numbers are model checkpoints used in experiments. *scGPT was trained on 6000 highly variable genes due to memory limitations with our setup. All other methods used all genes after standard filtering, which amounted to over 36,000 genes. Total GPU hours for each model can be computed by multiplying the number of GPUs with the train time.

| MODEL | FOUNDATION | # PARAMETERS | GPU | # GPUS | # TRAINING SAMPLES | # EPOCHS | TRAIN TIME (HRS) |
| --- | --- | --- | --- | --- | --- | --- | --- |
| SCGEN | NO | 59,967,903 | A5000 | 1 | 218,732 | 60 | ~ 1 |
| SCDIFFUSION | NO | 781,510,338 | A5000 | 1 | 218,732 | 1638 | ~ 78 |
| SCVI | NO | 23,580,274 | A5000 | 1 | 218,732 | 100 | ~ 1 |
| SCGPT* | YES | 51,342,849 | A100 40GB | 8 | 218,732 | 4 | ~ 9 |
| C2S (PYTHIA-160M) | NO | 162,322,944 | A100 40GB | 8 | 218,732 | 21 | ~ 29 |

#### Evaluation

- **Cell type generation** The performance of C2S was quantitatively evaluated using *k*-NN (K-Nearest Neighbors) accuracy and Gromov-Wasserstein distance, measured against the original ground truth cells. The *k*-NN classifier was fit on the test dataset, and we used the values *k* = 3, 5, 10, and 25. Gromov-Wasserstein distance was used to compare the similarity and structural alignment between generated outputs, which is not always captured by *k*-NN accuracy due to its local nature. In order to generate the cells, we used the same text prompts from our training dataset. A comparison of metrics when using unseen but similar prompts can be found in Table 10. Overall, our evaluation metrics give a well-rounded assessment of each model’s ability to generate biologically plausible cells.
- **Perturbed cell generation** All models were evaluated using Pearson R and Spearman R correlation metrics. The correlations are computed between mean gene expression values in the test dataset and the data generated by the model. We select the top 5000 highly variable genes from the training dataset for evaluation. From those highly variable genes, we select the top 20 differentially expressed genes between exposures of the same cell type and perturbation. We also compute the same metrics by differencing with the mean gene expression of the opposite exposure (chronic vs acute) in the training dataset. Out of 140 possible combinations of labels, 10 test cell type/perturbation combinations with different exposures from those seen during training were used to evaluate the models.

## Results

- **Cell type generation** As shown in Table 1, our C2S model outperformed all other methods. Notably, our C2S-trained Pythia-160m model outperforms the foundation model scGPT and also scDiffusion, which requires over 900 million parameters in its standard implementation for the immune tissue dataset. The performance of scVI and scGEN was similar due to their similar architectures, both being variational autoencoders. Our results show that causal language modeling can be adapted to single cell generation and outperform other commonly used generative architectures.
- **Perturbed cell generation** Table 2 shows our C2S model produces more accurate unseen perturbations than scGen and scGPT. These results potentially indicate that C2S models are able to leverage semantic representations of labels in a way non-language models like scGPT and scGen cannot.

### 4.3. Experiment 2: cell label prediction

#### Objective

The second experiment evaluate C2S’s capacity for cell label classification with complex, combinatorial cell labels involing multiple metadata information about cells. We hypothesize that the natural language generation capabilities of C2S from pretraining will transfer well to complex label classification compared to baseline methods.

#### Methodology

We compared C2S against several single-cell analysis methods, including two state-of-the-art (SOTA) single-cell foundational models, scGPT (Cui et al., 2023a) and Geneformer (Cui et al., 2022), as well as two non-single cell baseline classifiers. To evaluate the generalization ability of the model to out-of-distribution data, we utilized two bulk datasets - L1000 (Subramanian et al., 2017) and GTEx (Consortium, 2020) - as benchmark datasets, which follow a different data distribution compared to the single-cell datasets which comprised the pretraining datasets for each foundation model. We also evaluate all models on a human PBMC single-cell dataset treated with various combinatorial cytokine stimulations (Dong et al., 2023), as a third benchmark dataset.

For each dataset, metadata associated with the cell - cell type, tissue, drug perturabtions, dosage information, stimulations, etc. -was gathered to create information-rich multipart labels for each cell/bulk sample in each dataset. Predicting combinations of labels is expected to be a challenging task, particular in a limited data scenario. We constrained the number of training samples for each dataset to roughly give a few samples per combinatorial class, resulting in a dataset where there are few examples per class. A comparison of compute resources for each of the foundation models can be found in Table 5.

#### Evaluation

We report the classification accuracy and Area Under the Receiver Operating Characteristic Curve (AUROC) metrics across 3 experiment repeats on each dataset, as reported in Table 3. Exact combinatorial label matching was calculated for each dataset, in addition to another set of metrics giving models partial credit for getting individual parts of the combinatorial label correct.

#### Results

As indicated in Table 3, C2S outperformed baseline methods in both exact and partial label accuracy, demonstrating its capability in complex label prediction through natural language. The experiment particularly highlights C2S’s adaptibility to out-of-distribution data, which was not present anywhere in its training distribution, while also learning to predict complex multi-part labels. We additionally provide attention visualizations for highly-attended to genes in Section F of the Appendix, giving insight into which genes receive high attention by the LLM for predictions of specific drug compounds and cell lines.

### Experiment 3: abstract summary generation

#### Objective

We demonstrate the ability for C2S models to generate meaningful text carrying biological insight given a single cell sentence. Additionally, we show that C2S performance cannot be achieved by large language models without fine-tuning despite their extensive pretraining.

#### Methodology

We use GPT-2 small fine-tuned on cell sentences truncated to the top-100 most expressed genes from the multi-tissue dataset (see Appendix 4.1). At inference, we prompt C2S models with a natural language prompt and a single truncated cell sentence sampled from one of 19 held out evaluation studies. We generate abstract summaries with 30 cell sentences sampled from each evaluation study. This approach allows for a direct comparison of the generated abstracts with the ground-truth abstracts from associated evaluation publications. We benchmark against large language models in a 10-shot prompting setting. Each baseline model is prompted with a cell sentence and tasked with producing biological insights from the cell as an abstract summary (see the full prompt in Appendix E.1). We compare our approach against OpenAI’s GPT-3.5-Turbo-1106, as well as Mistral- 7B-Instruct (Jiang et al., 2023) and Mixtral-8×7B-Instruct (Jiang et al., 2024) quantized to 4-bit precision with AWQ (Lin et al., 2023). We also benchmark the performance of GPT-2 small and large pretrained checkpoints without C2S fine-tuning. The pretrained GPT-2 models are prompted in a zero-shot fashion due to their small contexts—the models are unable to accommodate the 4,000 tokens required for the 10-shot prompting setting (more details about the evaluation can be found in Appendix E).

#### Evaluation

Generated summaries are embedded and evaluated for pairwise similarity relative to the ground-truth abstracts (Xiao et al., 2023). We compute the mean pairwise cosine similarity between evaluation studies to test whether generated abstracts are significantly correlated to their ground-truths, more so than to other original abstracts. We apply the T and KS test on the mean cosine similari-ties. We also compute embedding distribution correlations (Pearson’s, Spearman’s) and distances (MMD, Wasserstein) between generated and original abstracts.

#### Results

We find that C2S was more adept at generating abstracts that closely align with the ground truth, compared to baseline large language models. Table 4 shows that only C2S is able to generate differentiated abstracts given a prompt cell sentence and large context. Additionally, C2S-generated summaries lie closer to the held out study abstracts in embedding space by a 50% margin on MMD (see Table 4). This experiment brings empirical evidence for the utility of cell sentences to derive biological insights from single-cell data. C2S learns to associate relevant language (e.g., tissue type, condition) to complex gene sequences, going beyond capabilities acquired during pretraining. We present a qualitative comparison of generated abstract summaries in Appendix Figure 12. Further examples of generated abstracts compared to their original counterparts are shown in Appendix G.1.

## 5. Discussion

Cell2Sentence is a novel approach for training large language models using single-cell transcriptomics data, converting gene expression profiles into sequences of text called *cell sentences*. This method involves ranking gene names by their expression levels to create a reversible encoding of biological data with minimal loss of information. Language models fine-tuned on these cell sentences outperform other foundation models such as Geneformer and scGPT on embedding tasks and generative tasks. Cell sentences, which can be integrated with textual annotations, are versatile for generation and summarization tasks, benefiting from natural language pretraining. We show pretrained language models trained on combinations of text and cell sentences leads to nascent capabilities of drawing insight from data not inherent to large language models such as GPT-3.5 and Mixtral. We leave the possibility of exploring the emergent properties of large language models with model and data scale to future work.

## Acknowledgements

We acknowledge the support of the National Institutes of Health R35 1R35GM143072-01 and R01 3R01AI157488-03S1 (to David van Dijk). Amin Karbasi acknowledges funding in direct support of this work from NSF (IIS-1845032) and the AI Institute for Learning-Enabled Optimization at Scale (TILOS).

## Impact Statement

This paper presents work whose goal is to advance the field of Machine Learning. There are many potential societal consequences of Cell2Sentence, particularly in the field of biomedical research. We hope that pretrained language models fine-tuned on single cell sentences will aid biological data generation and processing, uncovering valuable insights through a natural language perspective.

## A. Method Details

We further evaluate the robustness of the Cell2Sentence transformation across 127 single-cell datasets retrieved from (Megill et al., 2021). Among those are the 99 human single-cell datasets introduced in Section 4, as well as 28 mouse single-cell datasets. We quantify the reconstruction performance using a linear regression between log-rank and normalized expression on Pearson’s R, Spearman’s R and R-squared coefficients (see Figure 6). We also visualize the linear linear between log-rank and normalized expression across 12 human datasets in Figure 7. Interestingly, emerging patterns seem to hold across data samples within tissues (e.g., kidney and blood tissue samples achieve relatively similar reconstruction performance).

## B. Experimental Details

### B.1. Training Data Augmentation

We augment the original abstracts of studies in the multi-tissue dataset and generate 5,000 summaries for each abstract with GPT-3.5-Turbo-1106, yielding 495,000 unique abstract summaries (Eldan & Li, 2023; Taori et al., 2023b). The prompt employed to generate the abstracts is presented in Figure 8.

**Figure 8.**
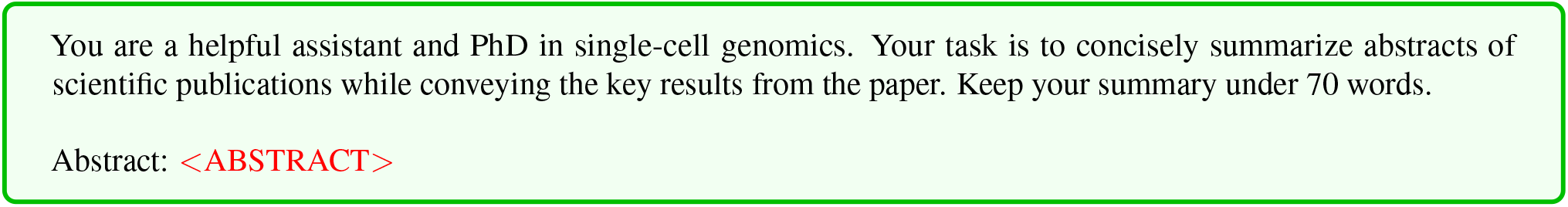
Prompt employed to generate synthetic abstract summaries using GPT-3.5-Turbo-1106.

### B.2. Evaluation Datasets

#### Cell label prediction

Our evaluation benchmarks model performance across three distinct datasets, incorporating both single-cell and bulk data. Each dataset introduces unique challenges for label classification, stemming from their combinatorial label structures and the presence of out-of-distribution samples:

1. Human PBMC single-cell (Dong et al., 2023): This dataset comprises 250 PBMC cells distributed among 7 cell types, incorporating 10 different stimulations and distinguishing between acute and chronic stimulations. The evaluation set encompasses approximately 120 unique label combinations, challenging the model’s ability to navigate sparse label spaces.

2. L1000 (Subramanian et al., 2017): Comprising bulk RNA sequencing data, the L1000 dataset is considered out-of-distribution for our task. It features 2,000 bulk samples, with labels denoting the cell line, drug compound (with 20 possible values), dosage, and perturbation time, leading to a total of 4,000 potential label combinations. This dataset serves to evaluate the model’s generalization capabilities across a broad spectrum of biological conditions and experimental interventions.

3. GTEx (Consortium, 2020): Similarly utilizing bulk RNA sequencing data, the GTEx dataset includes 1,000 bulk samples. Labels combine patient age, death condition (such as ventilator use or sudden death), and tissue type, amounting to around 1,000 distinct label combinations. Through this dataset, the model’s ability to deduce complex biological states from significantly diverse data compared to single-cell observations is scrutinized.

### B.3. Cell Generation

For cell type generation, we train the pretrained human scGPT model with sequence lengths of 1200 nonzero genes as described in (Cui et al., 2023a) for 1 epoch. In order for the model to generate zero-expressed genes, we train for 3 more epochs on randomly sampled collections of 2000 genes including zero-expressed genes with the same masking ratios used during the first phase of finetuning. The reported results show the model trained after its first epoch since the model had converged and stayed at the same loss for the remaining epochs. Even on a p4d.24xlarge AWS instance with 8 A100 40GB GPUs, half-precision, and flash attention 2, we found it difficult to fit longer sequences without memory issues. Although Pythia-160m has approximately 3 times as many parameters as scGPT, it was able train on the same instance 4 times faster per epoch despite having 4.5 times more tokens per sample.

The training procedure is simplified for perturbed cell generation since we restrict to 5000 highly variable genes in our evaluations. We train scGPT directly on these 5000 highly variable genes starting from the pretrained human model with mask ratios of .25, .50, and .75. The best model checkpoint is used for evaluation.

To generate cells at inference, we start with a fixed number of randomly sampled genes from a randomly sampled cell in our dataset. In the case of cell type generation, the cell is sampled from the training dataset. For perturbed cell generation, the cell is sampled from the test dataset. A cell is generated autoregressively at inference using the previously generated genes as context until all genes are generated. The number of genes used as context is 1000 for cell type generation and 2500 for perturbed cell generation. The same number of genes are generated during each forward pass. Labels are embedded and added to all other tokens, so the model receives two sets of conditional signals during inference.

## C. Training Details

We provide further details on our training configuration for the pretraining for experimental results and fine-tuning settings. We refer to natural language as “NL” and Cell2Sentence as “C2S”, and refer to a model that has been solely trained on cell sentences as “C2S”, in contrast to a pretrained model fine-tuned on cell sentences which we refer to as “NL + C2S”. We use sequence lengths of 1024 for GPT-2 and 9200 for Pythia-160m and train on all tokens. We use the AdamW optimizer (Loshchilov & Hutter, 2017) and flash attention (Dao et al., 2022; Dao, 2023). We find that C2S models largely benefit from natural language pretraining as opposed to starting training from randomly initialized weights as explained below (see Appendix C.1).

### C.0.1. PRETRAINING

The GPT-2 small model is initialized with 12 layers and 768 hidden dimensions, and the medium model with 24 layers and 1024 hidden dimensions, as detailed in (Radford et al., 2019). We employ a learning rate of 6 *×* 10^−4^ with a cosine scheduler and 1% warmup ratio. For the GPT-2 medium model, we accumulate gradients over 16 steps. The effective batch sizes for the small and medium models are of 10 and 48 examples. Each model is trained using a single A5000 GPU over two days. Model weights are randomly initialized using a Xavier normal distribution (Glorot & Bengio, 2010).

We train a Byte Pair Encoding (BPE) tokenizer (Sennrich et al., 2015) on the full cell sentence dataset, including NL prompts and cell type labels, yielding a vocabulary of 9,609 tokens. The training set contains approximately 30 million tokens, averaging 740 tokens per example. Due to the smaller embedding space, the initialized models contain slightly fewer parameters than their counterparts pretrained on a vocabulary of 50,257 tokens (93M for the small model and 313M for medium model). The resulting corpus exhibits sparse NL tokens due to short and repetitive prompts. Despite instruction corpora being traditionally used to fine-tune pretrained models for question answering tasks, we adopt this setting during pretraining to mirror our fine-tuning setup described in Section C.0.2. We hypothesize that the semantic variability from prompting patterns might implicitly regularize token and positional embeddings, with natural language tokens acting as class tokens.

We emphasize that the loss is computed on both the prompt and the associated label (i.e. cell type). Not doing so would cause embeddings of the prompt tokens to remain random, impairing the capacity of the model to learn the conditional relations between prompt and label tokens. We evaluate the capacity of our model to generate valid genes and maintain an accurate sequence length (here, of 100 genes) and present the results in Table 11. We find that both pretrained models are able to generate sequences of 100 genes without significantly deviating from the mean. The models also both achieve over 97% and 96% accuracy in gene validity and uniqueness.

### C.0.2. FINE-TUNING

Models are initialized using pretrained weights retrieved from the Hugging Face model hub (HF Canonical Model Maintainers, 2022). We employ a cosine scheduler. On both models, we accumulate gradients over 16 steps and use batch sizes of eight examples (yielding an effective gradient update batch size of 128 examples). Each model is trained using a single A5000 GPU. While we experimented with applying efficient fine-tuning techniques (e.g. LoRA (Hu et al., 2022)), fully fine-tuned models outperformed alternatives in gene uniqueness and validity assessments. We notably found LoRA to yield highly variable generation patterns, with uniqueness of genes in generated sentences as low as 70%. Unlike for our pretraining setup, we apply the instruction fine-tuning task in a classical manner, computing the loss exclusively on labels. We use the pretrained GPT-2 tokenizer, which averages around 233 tokens per training samples (yielding a total of 9M training tokens).

Similarly to the process detailed in Section C.0.1, we examine the coherence of generated output using sequence length, as well as accuracy in gene validity and uniqueness. We find that the fine-tuned model outperform the pretrained models by generating genes with over 99% validity and 98% uniqueness on average (see Table 11). While both models achieve reliable performance by these standard metrics, we conclude that our fine-tuned models are consistently outperforming the pretrained models in generating real human genes, which are only rarely duplicated within cell sentences.

### C.0.3. COMPUTE RESOURCES

Comparisons of the amount of compute required to train and fine-tune models for our experiments are found in Tables 5 and 6. When adjusting for the number of parameters and number of epochs, C2S models compare favorably to other foundation models. However, wholly fair comparisons are challenging due to the uniqueness of C2S models’ integrated text inputs and its leveraging of highly optimized open source libraries.

### C.1. Comparing fine-tuning and pretraining

We test whether using pretrained GPT-2 weights yields performance improvements over only training from randomly initialized weights on cell sentences. We therefore train two GPT-2 small models on a random subset of 49,920 cell sentences spanning 17 cell types from the immune tissue dataset. To evaluate the ability of our trained models to generate realistic cells, we consider the average generated cell of each of the 17 cell types in our immune tissue dataset, and compare it with the average real cell of each cell type in Table 7. Across 17 different cell types, generated cells from fine-tuned models show high correlation with real cells, capturing over 94% of the variation in the expression of an average cell. We note that initializing a model with a pretrained language model outperforms training from scratch, indicating that there is mutual information which allows a model to better understand cell sentence generation.

**Table 7.** Correlation metrics for averaged generated cells per cell type against original expression values. The “random” model was trained from scratch on cell sentences. The “pretrained” model was fine-tuned on cell sentences. Correlation metrics are computed on individual cell types and then averaged.

| MODEL | PEARSON R | $R^2$ |
| --- | --- | --- |
| GPT-2 SMALL, RANDOM | 0.947 | 0.873 |
| GPT-2 SMALL, PRETRAINED | **0.984** | **0.949** |

We also trained an additional two GPT-2 medium models to assess the performance improvements brought by scaling the parameter count. Table 8 show the *k*-NN accuracy of each model for different number of neighbors. The cell type of cells generated by C2S models are accurately classified with a *k*-NN classifier, achieving a peak of accuracy of 54%. Table 9 underscores the necessity of NL pretraining for accurate cell type identification. A significant performance decline is observed when using models that have not undergone NL pretraining, thereby confirming that the models are not merely memorizing the conditioning text. Despite 1) the limited scope of natural language text in our training prompts relative to the pretraining corpus, and 2) permitting the models to train on these natural language prompts, models without NL pretraining failed to acquire meaningful natural language embeddings. Furthermore, a modest performance increment is observed as the scale of pretrained models increases.

**Table 8.** k-NN classification accuracy results against ground truth data. “NL + C2S” means pretrained on natural language and then trained on cell sentences. “C2S” means no pretraining and just trained on cell sentences. *k*-NN classifier is fitted on ground truth cell sentences and used to predict the cell type label of generated cell sentences from different trained models. *k*-NN classification is done both in cell sentence space using Levenshtein distance (Lev.), as well as after converting back to expression vectors (Expr.). “Real cells” indicates *k*-NN classification fit on ground truth cell sentences and used to predict a separate sample of ground truth cell sentences.

| MODEL | $k=5$ | | $k=10$ | | $k=25$ | | $k=50$ | |
| --- | --- | --- | --- | --- | --- | --- | --- | --- |
|  | EXPR. | LEV. | EXPR. | LEV. | EXPR. | LEV. | EXPR. | LEV. |
| GPT-2 SMALL (C2S) | 24.56 | 16.02 | 23.79 | 17.72 | 23.51 | 19.17 | 22.84 | 19.76 |
| GPT-2 SMALL (NL + C2S) | 52.55 | 37.63 | 52.19 | 41.20 | 51.42 | 43.42 | 49.51 | 44.44 |
| GPT-2 MEDIUM (C2S) | 26.36 | 17.58 | 25.48 | 19.18 | 24.55 | 20.67 | 23.44 | 21.32 |
| GPT-2 MEDIUM (NL + C2S) | **54.67** | **38.60** | **54.52** | **41.34** | **53.27** | **43.93** | **51.80** | **44.73** |

**Table 9.** Quantification of autoregressive cell type prediction on unseen cells. “NL + C2S” means pretrained on natural language and then trained on cell sentences. “C2S” means no pretraining and just trained on cell sentences. These results show test accuracy significantly improves with NL pretraining. The scores are computed on unseen immune tissue test data and weighted by the distribution of labels.

| MODEL | ACCURACY | F1 | PRECISION | RECALL |
| --- | --- | --- | --- | --- |
| GPT-2 SMALL (C2S) | 29.33 | 17.46 | 13.27 | 29.33 |
| GPT-2 SMALL (NL + C2S) | 69.95 | 68.56 | 69.44 | 69.95 |
| GPT-2 MEDIUM (C2S) | 29.17 | 17.41 | 13.35 | 19.17 |
| GPT-2 MEDIUM (NL + C2S) | **74.26** | **73.69** | **74.16** | **74.26** |

**Table 10.** Comparison of *k*-NN performance on cell type generation with the C2S (Pythia-160m) model when using seen vs unseen prompts. This is the same setup as the results reported in Table 1, except the unseen prompts are new prompts generated by GPT-4 which were not used during training. The new prompts are similar in semantic meaning to the original training prompts. Our results show there is some robustness to variability in prompts at our model scale, but there is an observable reduced performance. We hypothesize that larger and improved models would suffer less from changes in training prompts.

| PROMPTS | $k$-NN (↑) | | | |
| --- | --- | --- | --- | --- |
|  | 3 | 5 | 10 | 25 |
| SEEN | $0.2588 \pm 0.0061$ | $0.2565 \pm 0.0060$ | $0.2746 \pm 0.0073$ | $0.2715 \pm 0.0070$ |
| UNSEEN | $0.1983 \pm 0.0050$ | $0.1949 \pm 0.0067$ | $0.2083 \pm 0.0065$ | $0.2022 \pm 0.0056$ |

**Table 11.** Quality of generated outputs. “NL + C2S” means pretrained on natural language and then trained on cell sentences. “C2S” means no pretraining and just trained on cell sentences. This table shows 1. models trained using Cell2Sentence are able to generate real genes with few duplicates and invalid genes and 2. models pretrained with natural language generate more accurately. The metrics are computed across all 35 cell types seen during training with 500 cells generated per cell type and then averaged across all generated cells (top 100 genes) from the immune tissue dataset. The valid genes percentage shows the number of genes generated that are real genes including duplicates. The generated length is the number of genes generated regardless of their validity. The unique gene ratio is the ratio of unique valid genes to the generated length.

| MODEL | GEN. LENGTH | VALID GENES % | UNIQUE GENES % |
| --- | --- | --- | --- |
| GPT-2 SMALL (C2S) | 101.54 | 97.84 | 97.31 |
| GPT-2 SMALL (NL + C2S) | 99.84 | 99.60 | 98.88 |
| GPT-2 MEDIUM (C2S) | 100.62 | 97.51 | 96.69 |
| GPT-2 MEDIUM (NL + C2S) | 99.80 | **99.70** | **99.47** |

## D. Inference Details

At inference, we follow the training procedure highlighted in Section C. We set the hyper-parameters top p = 0.9 and temperature = 0.7 to promote diversity in cell generation. For cell type generation and unconditional cell generation used in our experiments (see Section 4), we randomly sample prompt templates as input, inserting the cell type or sentence where needed for conditional generation. For autoregressive cell type prediction or abstract generation, we randomly sample templates as in training.

All outputs are generated until an end-of-sequence (EOS) token is predicted. Post-generation, gene and cell type extraction is done using regex to remove prompts. For evaluation, we retain invalid genes and average ranks of duplicate genes, rearranging sequences as needed. When reverting back to expression values, invalid genes are ignored, but the rank values are preserved, e.g. if an invalid gene appears in position 3 and a valid gene appears in position 4, the invalid gene is ignored, but the valid gene retains a rank of 4.

Utilizing pretrained LLMs offers the advantage of using highly optimized, open-source libraries for inference. Similarly to the training setup, we make use of flash attention (Dao et al., 2022; Dao, 2023) and batched inference to accelerate generation. Inference for GPT-2 medium (345M parameters) can be done on a single A5000 GPU with 24GB of VRAM and a batch size of 100 without running out of memory. For GPT-2 small (117M parameters), the batch size can be increased to 250. We did not determine the exact maximum batch sizes, so these values can likely be increased further. On average, the number of tokens in the prompts and top 100 genes combined was around 350. For example, the GPT-2 small model takes approximate 20 minutes to generate 500 cells from each of the 35 cell types found in the immune tissue dataset. Model quantization was not required but may be useful for future experiments with larger models.

## E. Evaluation

### E.1. In-Context Learning Abstract Generation

We evaluate the capacity of closed- and open-source LLMs to generate biological insights from cell sentences in a 10-shot prompting setting. Each model is provided with a sequence of pairs of cell sentences and abstract summaries. We evaluate GPT-3.5-Turbo-1106 with the prompt presented in Figure 11. We use an identical prompt for Mixtral-8×7B-Instruct and Mistral-7B-Instruct, while removing the instruction to return JSON format as this is not known to be a feature of these models. This 10-shot prompting approach yields a context length of around 4000 tokens. As GPT-2 small and GPT-2 large are limited to 1024 in-context tokens, we use the same prompt as for the C2S models, in which we only provide a minimal instruction in natural language paired with a cell sentence.

## F. Attention Visualizations

We provide attention visualizations for highly-attended to genes in drug compound (Figure 9) and cell line (Figure 10) predictions. Attention coefficients corresponding to input genes are first aggregated for drug compound and cell line label predictions across all test samples in the L1000 dataset. We extract attention coefficients of the last hidden layer of our finetuned GPT-2 model, and average attention coefficients across different attention heads. Gene attention coefficients are then averaged for each unique drug compound (Figure 9, y-axis) or cell line (Figure 10, y-axis) label, and the top 50 highly-attended to genes are visualized. The biclustered heatmaps reveal differential attention of genes for the prediction of different labels, giving insight of which input genes in the cell sentences the model is attending to for prediction of different drug compounds and cell lines.

**Figure 9.**
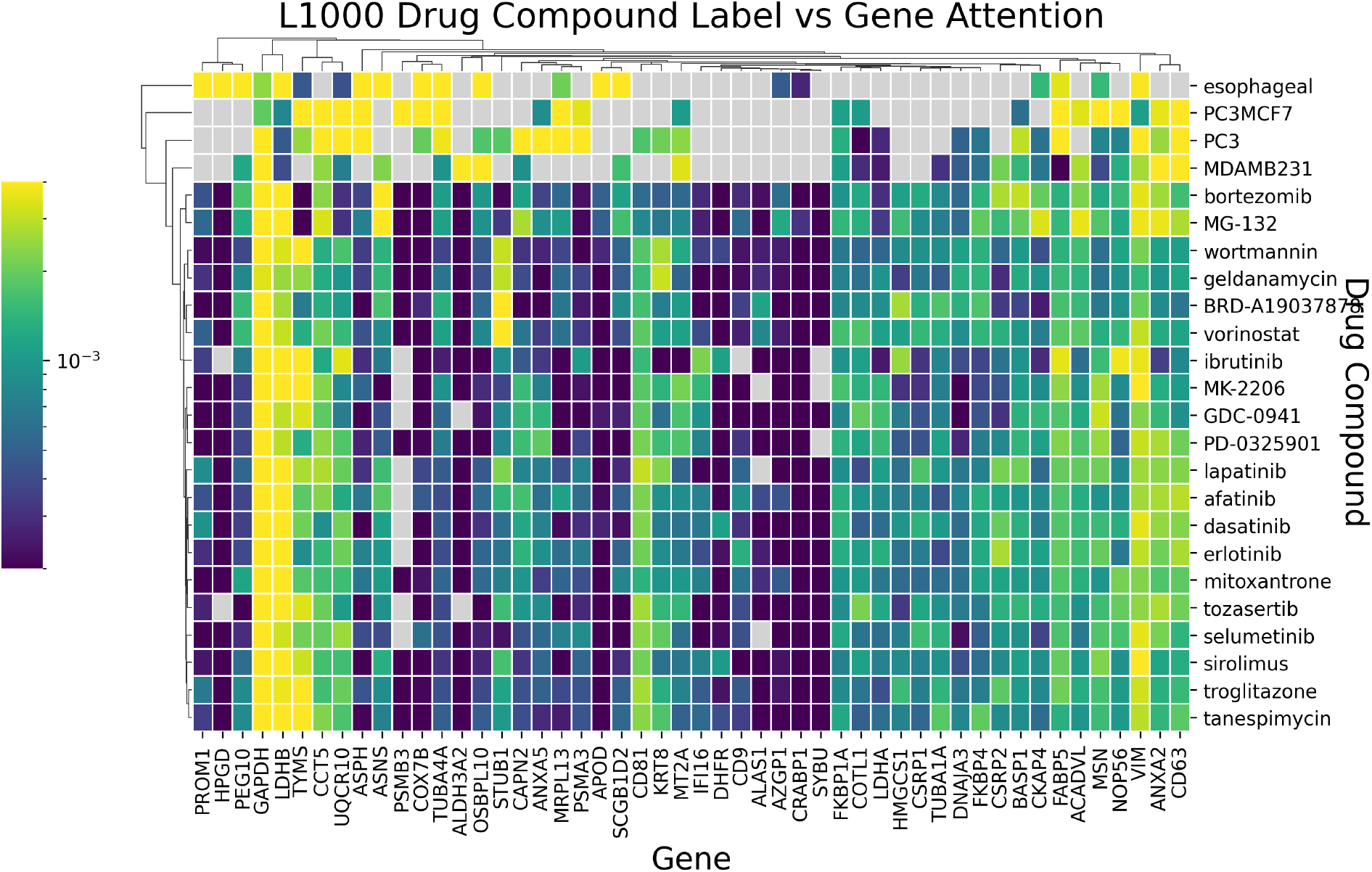
Attention heatmap visualization for drug compound prediction in the L1000 dataset.

**Figure 10.**
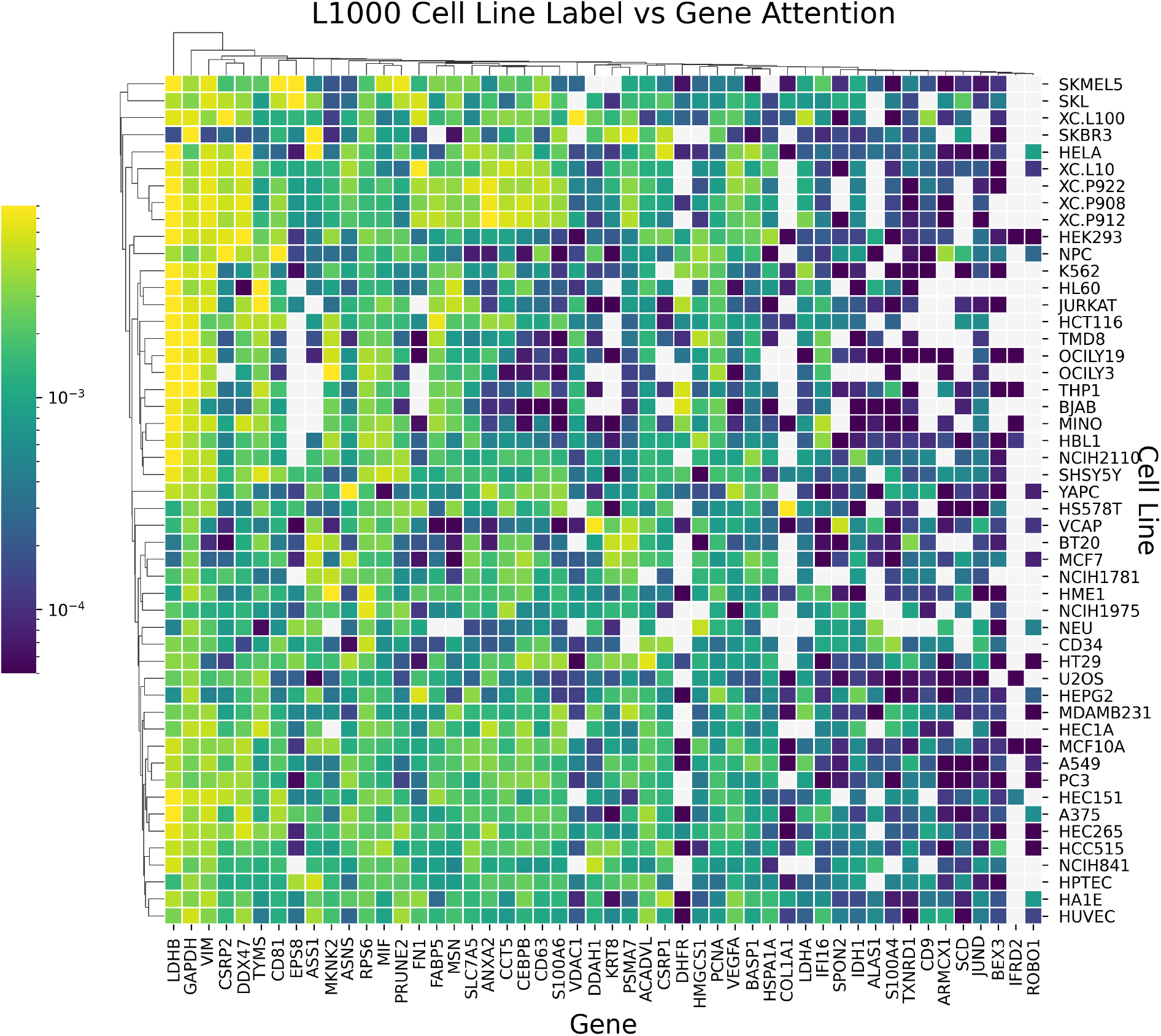
Attention heatmap visualization for cell line prediction in the L1000 dataset.

**Figure 11.**
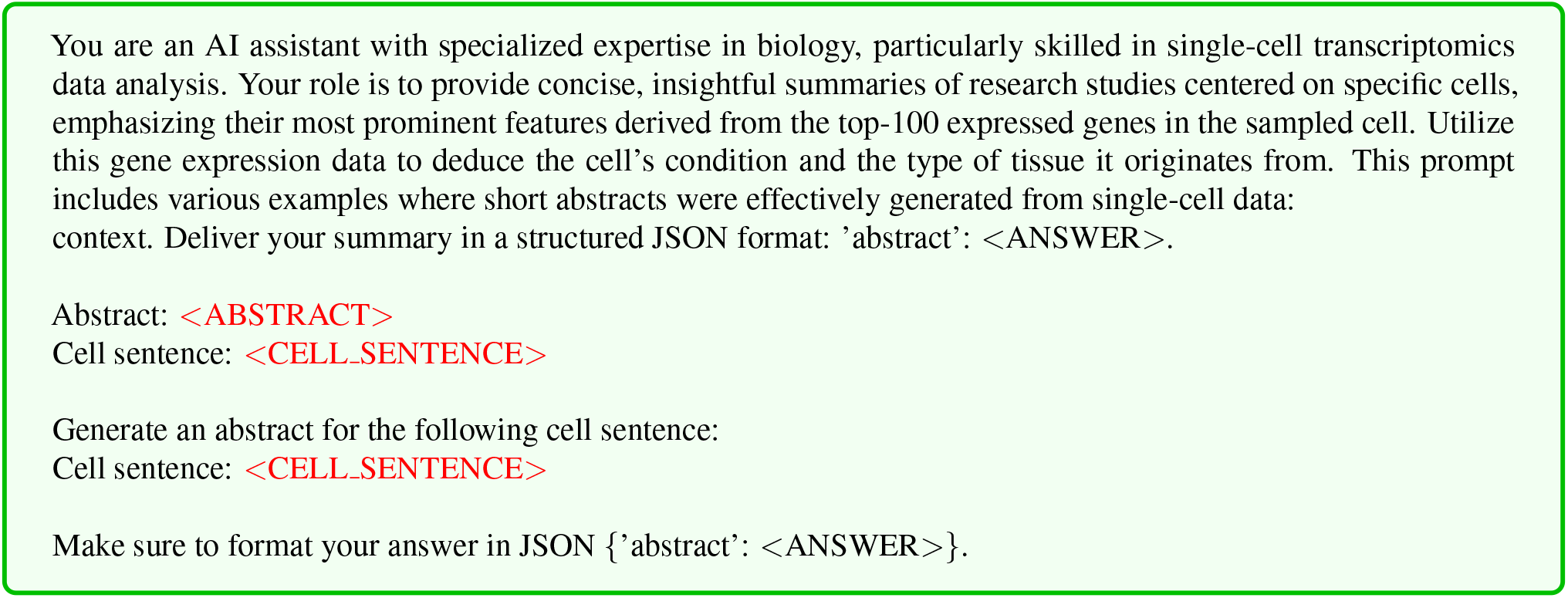
In-context learning prompt employed to generate abstract summaries from cell sentences with GPT-3.5-Turbo-1106.

**Figure 12.**
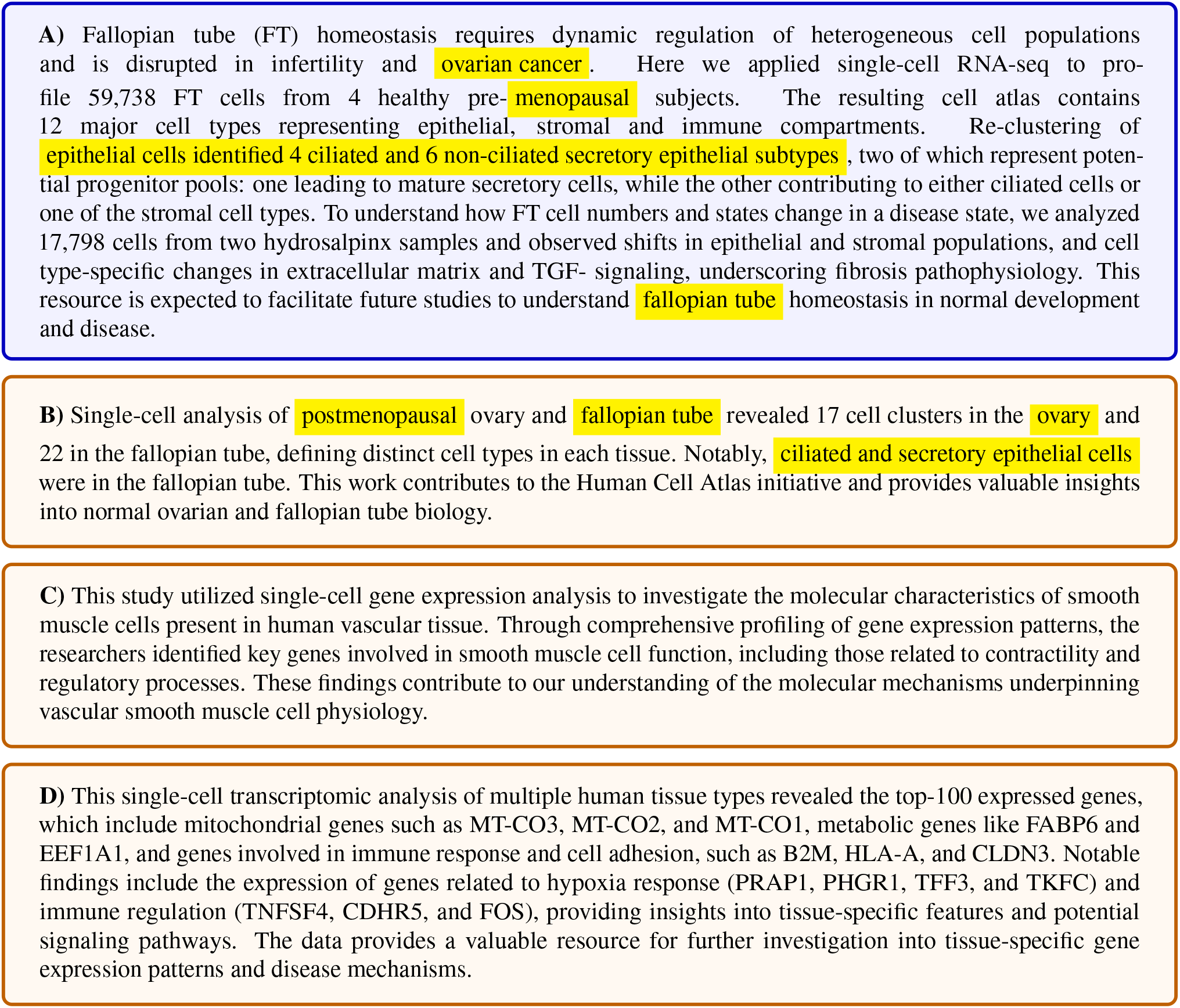
Comparison between an original study abstract (A), and abstract summaries generated with GPT-2 small fine-tuned with C2S (B), GPT-3.5-Turbo-1106 (C), and Mixtral-8×7B-Instruct (D). Only the C2S-generated abstract is relevant to the ground-truth study abstract, despite all summaries being semantically correct. Note that this study was held out from the training set.

## G. Generated abstracts

### G.1. Examples of generated abstracts

The following examples of generated abstracts show C2S models can generate natural language insights from single cell sentences not necessarily contained in the original abstract of the corresponding study.

#### Original study abstract

Although most SARS-CoV-2-infected individuals experience mild coronavirus disease 2019 (COVID-19), some patients suffer from severe COVID-19, which is accompanied by acute respiratory distress syndrome and systemic inflammation. To identify factors driving severe progression of COVID-19, we performed single-cell RNA-seq using peripheral blood mononuclear cells (PBMCs) obtained from healthy donors, patients with mild or severe COVID-19, and patients with severe influenza. Patients with COVID-19 exhibited hyper-inflammatory signatures across all types of cells among PBMCs, particularly up-regulation of the TNF/IL-1*β*-driven inflammatory response as compared to severe influenza. In classical monocytes from patients with severe COVID-19, type I IFN response co-existed with the TNF/IL-1*β*-driven inflammation, and this was not seen in patients with milder COVID-19. Interestingly, we documented type I IFN-driven inflammatory features in patients with severe influenza as well. Based on this, we propose that the type I IFN response plays a pivotal role in exacerbating inflammation in severe COVID-19. (Lee et al., 2020)

#### C2S generated abstract

In patients with critical COVID-19, 19% had type I interferon autoantibodies, compared to 6% in severe cases and none in moderate cases. Analysis of immune cells from patients revealed reduced type I interferon response in myeloid cells, particularly in dendritic cells with autoantibodies. Monocytes from critically ill patients showed elevated expression of the inhibitory receptor LAIR1, which correlated with reduced interferon response. These findings suggest a unified mechanism involving interferon suppression in sever COVID-19.

#### Original study abstract

Fallopian tube (FT) homeostasis requires dynamic regulation of heterogeneous cell populations and is disrupted in infertility and ovarian cancer. Here we applied single-cell RNA-seq to profile 59,738 FT cells from 4 healthy pre-menopausal subjects. The resulting cell atlas contains 12 major cell types representing epithelial, stromal and immune compartments. Re-clustering of epithelial cells identified 4 ciliated and 6 non-ciliated secretory epithelial subtypes, two of which represent potential progenitor pools: one leading to mature secretory cells, while the other contributing to either ciliated cells or one of the stromal cell types. To understand how FT cell numbers and states change in a disease state, we analyzed 17,798 cells from two hydrosalpinx samples and observed shifts in epithelial and stromal populations, and cell type-specific changes in extracellular matrix and TGF-signaling, underscoring fibrosis pathophysiology. This resource is expected to facilitate future studies to understand fallopian tube homeostasis in normal development and disease. (Ulrich et al., 2021)

#### C2S generated abstract

This study provides single-cell transcriptomic and regulatory profiles of postmenopausal ovary and fallopian tube. In the fallopian tube, 22 cell clusters integrated into 11 cell types, including ciliated and secretory epithelial cells. The ovary had 17 distinct cell clusters defining 6 major cell types. These findings contribute to the understanding of cellular composition in these tissues for future studies on reproductive health and diseases.

#### Original study abstract

Gastrointestinal microbiota and immune cells interact closely and display regional specificity; however, little is known about how these communities differ with location. Here, we simultaneously assess microbiota and single immune cells across the healthy, adult human colon, with paired characterization of immune cells in the mesenteric lymph nodes, to delineate colonic immune niches at steady state. We describe distinct helper T cell activation and migration profiles along the colon and characterize the transcriptional adaptation trajectory of regulatory T cells between lymphoid tissue and colon. Finally, we show increasing B cell accumulation, clonal expansion and mutational frequency from the cecum to the sigmoid colon and link this to the increasing number of reactive bacterial species. (James et al., 2020)

#### C2S generated abstract

In this study, 720,633 cells from the intestines of 71 individuals with Crohn’s disease were analyzed to understand cell-specific responses to inflammation. The findings showed distinct responses in immune, epithelial, and stromal cells between the small and large intestines, particularly related to genes associated with inflammatory bowel disease (IBD) risk. Additionally, potential regulators of fibrotic complications were identified, offering insights into potential therapeutic targets for Crohn’s disease. This work provides a roadmap for understanding the disease at a cellular level.

#### Original study abstract

During postnatal life, thymopoiesis depends on the continuous colonization of the thymus by bone-marrow-derived hematopoietic progenitors that migrate through the bloodstream. The current understanding of the nature of thymic immigrants is largely based on data from pre-clinical models. Here, we employed single-cell RNA sequencing (scRNA-seq) to examine the immature postnatal thymocyte population in humans. Integration of bone marrow and peripheral blood precursor datasets identified two putative thymus seeding progenitors that varied in expression of CD7; CD10; and the homing receptors CCR7, CCR9, and ITGB7. Whereas both precursors supported T cell development, only one contributed to intrathymic dendritic cell (DC) differentiation, predominantly of plasmacytoid dendritic cells. Trajectory inference delineated the transcriptional dynamics underlying early human T lineage development, enabling prediction of transcription factor (TF) modules that drive stage-specific steps of human T cell development. This comprehensive dataset defines the expression signature of immature human thymocytes and provides a resource for the further study of human thymopoiesis. (Lavaert et al., 2020)

#### C2S generated abstract

This study combined single-cell genomics techniques to reconstruct the developing human immune system across nine prenatal tissues. The findings revealed the timing of immune cell subset acquisition and maturation, as well as blood and immune cell development beyond primary hematopoietic organs. The study also characterized human prenatal B1 cells and shed light on the origin of unconventional T cells, providing valuable insights for cell engineering, regenerative medicine, and disease understanding.

#### Original study abstract

The retina is a specialized neural tissue that senses light and initiates image processing. Although the functional organization of specific retina cells has been well studied, the molecular profile of many cell types remains unclear in humans. To comprehensively profile the human retina, we performed single-cell RNA sequencing on 20,009 cells from three donors and compiled a reference transcriptome atlas. Using unsupervised clustering analysis, we identified 18 transcriptionally distinct cell populations representing all known neural retinal cells: rod photoreceptors, cone photoreceptors, Mü ller glia, bipolar cells, amacrine cells, retinal ganglion cells, horizontal cells, astrocytes, and microglia. Our data captured molecular profiles for healthy and putative early degenerating rod photoreceptors, and revealed the loss of MALAT1 expression with longer post-mortem time, which potentially suggested a novel role of MALAT1 in rod photoreceptor degeneration. We have demonstrated the use of this retina transcriptome atlas to benchmark pluripotent stem cell-derived cone photoreceptors and an adult Müller glia cell line. This work provides an important reference with unprecedented insights into the transcriptional landscape of human retinal cells, which is fundamental to understanding retinal biology and disease. (Lukowski et al., 2019)

#### C2S generated abstract

This study used single-cell RNA sequencing to analyze human retinal cells, identifying 58 cell types and mapping gene expression patterns linked to blinding diseases. They found differences in gene expression and proportions between foveal and peripheral retinal cells. Comparing human and macaque retinas, they found over 90% correspondence in cell types and expression of genes implicated in blinding diseases, validating the macaque as a model for studying blinding diseases.

#### Original study abstract

The liver is the largest solid organ in the body and is critical for metabolic and immune functions. However, little is known about the cells that make up the human liver and its immune microenvironment. Here we report a map of the cellular landscape of the human liver using single-cell RNA sequencing. We provide the transcriptional profiles of 8444 parenchymal and non-parenchymal cells obtained from the fractionation of fresh hepatic tissue from five human livers. Using gene expression patterns, flow cytometry, and immunohistochemical examinations, we identify 20 discrete cell populations of hepatocytes, endothelial cells, cholangiocytes, hepatic stellate cells, B cells, conventional and non-conventional T cells, NK-like cells, and distinct intrahepatic monocyte/macrophage populations. Together, our study presents a comprehensive view of the human liver at single-cell resolution that outlines the characteristics of resident cells in the liver, and in particular provides a map of the human hepatic immune microenvironment. (MacParland et al., 2018)

#### C2S generated abstract

This study used single-cell RNA sequencing and single nucleus RNA sequencing to comprehensively map the cell types in the human liver. The addition of snRNA-seq revealed new subtypes of hepatic stellate cells and cholangiocyte progenitors, while T and B lymphocytes and NK cells were only distinguishable using snATAC-seq. The study validated the spatial distribution of liver cell populations using spatial transcriptomics and immunohistochemistry. This work provides a high-resolution map of healthy human liver parenchymal cell populations.

